# Maternal provisioning of an obligate symbiont

**DOI:** 10.1101/2022.09.07.506999

**Authors:** Tyler J. Carrier, Lara Schmittmann, Sabrina Jung, Lucía Pita, Ute Hentschel

## Abstract

Vertical transmission of microbial symbionts is interpreted as all offspring within a clutch being provided a similar number of symbionts irrespective of reproductive output (fecundity). This interpretation, however, stems primarily from oviparous insects and, thus, has yet to consider other major reproductive strategies. We used the viviparous sponge *Halichondria panicea* and its obligate symbiont “*Candidatus* Halichondribacter symbioticus” to test the hypothesis that offspring receive quantitatively similar numbers of its obligate symbiont. This quantitative strategy of vertical transmission was not observed. Instead, we find that *H. panicea* has a maternal pool of ‘*Ca*. H. symbioticus’ that is partitioned proportionally to reproductive output and allometrically by offspring size. Moreover, ‘*Ca*. H. symbioticus’ could not be experimentally reduced in larvae by antibiotics, while the total bacterial community could be depleted. The ability to undergo metamorphosis was unaffected by this perturbation. Together, this demonstrates that the obligate symbiont ‘*Ca*. H. symbioticus’ is maternally provisioned and, thus, provides an additional strategy for how microbes can be vertically transmitted.

## INTRODUCTION

Animals from all major phylogenetic lineages use their reproductive machinery to vertically transmit microbes to the next generation [1–4]. A threshold in symbiont quantity is often required to ensure that these partnerships are maintained through the developmental stages [1–4]. This minimum can span six orders of magnitude (from 10s to >1 million) and can incur an inter-sibling variability of less than 20% [5–9]. The aphid *Uroleucon ambrosiae*, for example, provides ~10^3^ (± ~5%) cells of the nutritional mutualist *Buchnera* spp. to each egg, while the stinkbug *Megacopta punctatissima* produces one capsule with ~10^8^ cells of the obligate symbiont ‘*Candidatus* Ishikawaella capsulate’ for every 3 to 4 eggs [9, 10]. This interpretation of vertical transmission—that all offspring within a clutch of any fecundity are provided a similar number of symbionts [1–4]—stems primarily from oviparous insects [5, 8–10] and, thus, has yet to consider other major reproductive strategies.

One such strategy is viviparity. In this, fertilization occurs internally, early embryos are brooded within the maternal body, and late-stage offspring are then released into the environment. Viviparity is widespread amongst animals and is particularly common in marine sponges. Brooding provides the adult sponge with a developmental window to maternally transmit microbes that are enriched within the mesohyl [11–16]. This strategy is particularly evident in ‘low microbial abundance’ sponges, which inherit a select few dominant symbionts that are essential for maternal, and sometimes offspring, fitness [11, 17]. One such sponge is *Halichondria panicea*, a coastal species that inhabits large parts of the North Atlantic and has a bacterial community that is dominated (with a 90:1 cell count) by the obligate symbiont “*Candidatus* Halichondribacter symbioticus” [18–21].

We used the sponge *H. panicea* to test the hypothesis that offspring between clutches have quantitatively similar numbers of ‘*Ca*. H. symbioticus’ (*i.e*., following that of oviparous insects). This quantitative strategy of vertical transmission was not observed. Instead, we find that *H. panicea* has a maternal pool of ‘*Ca*. H. symbioticus’ that is partitioned proportionally to reproductive output and allometrically by offspring size. Moreover, ‘*Ca*. H. symbioticus’ could not be experimentally reduced in larvae by antibiotics, while the total bacterial community could be depleted. The frequency that larvae undergo metamorphosis was unaffected by this perturbation. Together, this demonstrates that ‘*Ca*. H. symbioticus’ is maternally provisioned and, thus, provides an additional strategy for how microbes can be vertically transmitted.

## RESULTS

### Reproductive output and symbiont quantification

We collected adult *H. panicea* from the Kiel Bight (Western Baltic Sea, Germany) during late spring and early summer to quantitatively estimate the reproductive output of *H. panicea* and ‘*Ca*. H. symbioticus’ per offspring [22, 23]. All sponges were transferred within one hour to flow-through aquaria at the GEOMAR Helmholtz Centre for Ocean Research (Kiel, Germany). Mesh traps were placed around individual sponges [16], and seawater was collected regularly from the surface of each adult [24]. Total larvae from each adult on each collection day were counted and transferred to separate 1.5 mL Eppendorf tubes, of which were later pooled by adult sponge to quantify 16S rDNA copies for total bacteria and ‘*Ca*. H. symbioticus’ as well as the host cells (as estimated by the 18S rDNA gene) by quantitative polymerase chain reaction (qPCR).

Despite nearly three orders of magnitude difference between individuals (from 1 to 964 larvae), the reproductive output by this population was consistent across years (oneway ANOVA, F = 1.122, P = 0.332) (Fig 1A). Individuals that were observed to release larvae over a longer duration tended to have a greater reproductive output (F_1,40_ = 50.46, p < 0.0001; R^2^ = 0.558) (Fig S1A), with there being a high degree of variation between individuals and days (Fig S1B). Moreover, the reproductive output by this population increased gradually with sea surface temperature (Fig S1C) from mid-May (day number ~140) until early-June (day number ~160), after which there was a steady decline (SS: 1.129, R^2^ = 0.753, df = 21) (Fig 1B). This relationship between day number and reproductive output was not a biproduct of our sampling effort. Reproductive output increased linearly with the total number of sponges that were sampled on a given day (Fig S1D; F_1,23_ = 6.543, p = 0.018; R^2^ = 0.222) and with the total number of sponges that were sampled (Fig S1E; F_1,23_ = 16.21, p = 0.0005; R^2^ = 0.413).

**Fig 1:**
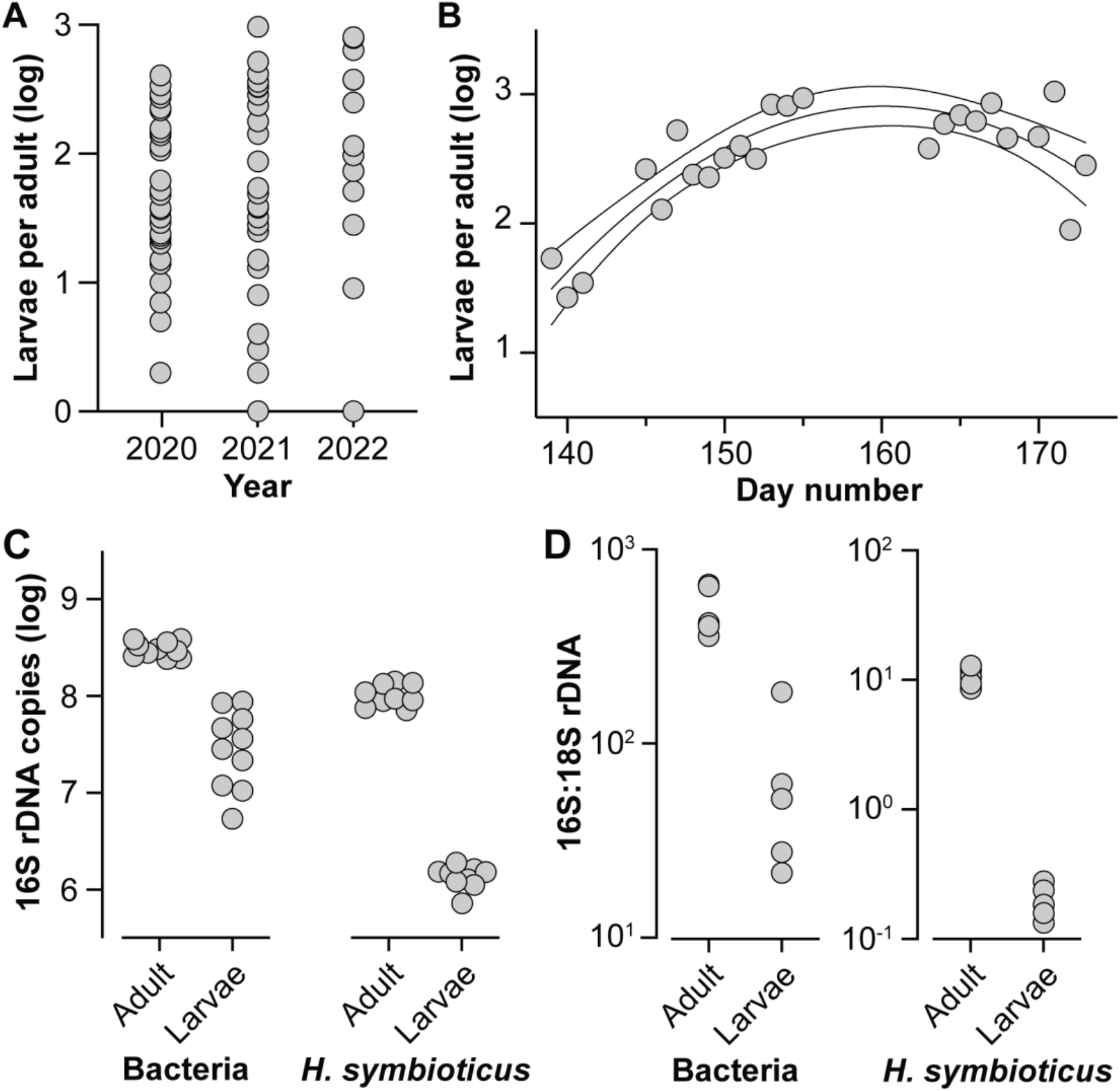
Reproductive output and symbiont transmission by *Halichondria panicea*. (A) Clutch size of adult *H. panicea* (n = 73) was consistent in three consecutive years (ANOVA, F = 1.122, P = 0.332). (B) The reproductive output of *H. panicea* was dynamic over the spawning season, such that there was a gradual increase in larval output from mid-May (day number 140) until early-June (day number 160) and a decline thereafter (SS: 1.129, R^2^ = 0.753, df = 21; average ± 95% confidence intervals). (C) Estimated number of 16S rDNA copies for total bacteria (left) and ‘*Candidatus* Halichondribacter symbioticus” (right) in adults and their clutch (n = 10, all from 2020). In both cases, the total 16S rDNA copies were significantly less in larvae than in adults (paired t-test, p < 0.0001 for both). (D) Estimated ratio of 16S to 18S rDNA copies for total bacteria (left) and “*Ca*. H. symbioticus’ (right) in adults and their clutch (n = 5, all from 2020). In both cases, larvae had less bacteria and ‘*Ca*. H. symbioticus’ per host cell than their respective adult (paired t-test, p < 0.0005 for both). All data was log transformed for normalization.

The estimated number of 16S rDNA copies per ng of DNA for total bacteria was significantly (7.9x) higher in adults than in their larvae (paired t-test, p < 0.0001) (Fig 1C). This pattern was also observed for ‘*Ca*. H. symbioticus,’ whereby adults had 81.3x more ‘*Ca*. H. symbioticus’ than larvae (paired t-test, p < 0.0001) (Fig 1C). Total 16S rDNA copies for all bacteria and for ‘*Ca*. H. symbioticus’ outnumbered the 18S rDNA copies of *H. panicea* for both adults and larvae (Fig 1D). Moreover, larvae had 7.2x less bacteria (paired t-test, p < 0.0001) and 65.5x less ‘Ca. H. symbioticus’ (paired t-test, p = 0.0002) per 18S rDNA copy than their respective adult (Fig 1D).

### *Maternal provisioning of* ‘Ca. *H. symbioticus’*

We used these quantitative estimates of reproductive output for *H. panicea*, host cells (a proxy for offspring size), total bacteria, and ‘*Ca*. H. symbioticus’ to test whether offspring between clutches have quantitatively similar numbers of symbionts (Fig 1, S1). We did not observe that offspring between different *H. panicea* individuals had similar numbers of ‘*Ca*. H. symbioticus.’ Instead, we observed a trade-off between fecundity and ‘*Ca*. H. symbioticus’ per larva (F_1,8_ = 32.20, p = 0.0005; R^2^ = 0.801), such that larvae from more fecund adults had less ‘*Ca*. H. symbioticus’ on average than larvae from less fecund adults (Fig 2A). This pattern, however, was not observed for total bacteria (F_1,8_ = 3.53, p = 0.097; R^2^ = 0.306) (Fig S2A). Moreover, we observed a trade-off between reproductive output, larval size, and ‘*Ca*. H. symbioticus,’ whereby larvae from more fecund sponges had less copies of the 18S rDNA gene and ‘*Ca*. H. symbioticus’ per larva than that from less fecund sponges (larval size vs reproductive output: F_1,3_ = 27.21, p = 0.014, R^2^ = 0.901; larval size vs ‘*Ca*. H. symbioticus’: F_1,3_ = 12.17, p = 0.039, R^2^ = 0.802) (Fig 2D, E). This pattern, again, was not observed for total bacteria (larval size vs reproductive output: F_1,3_ = 27.21, p = 0.097, R^2^ = 0.656; larval size vs total bacteria: F_1,7_ = 4.253, p = 0.078, R^2^ = 0.378) (Fig S2D, E).

**Fig 2:**
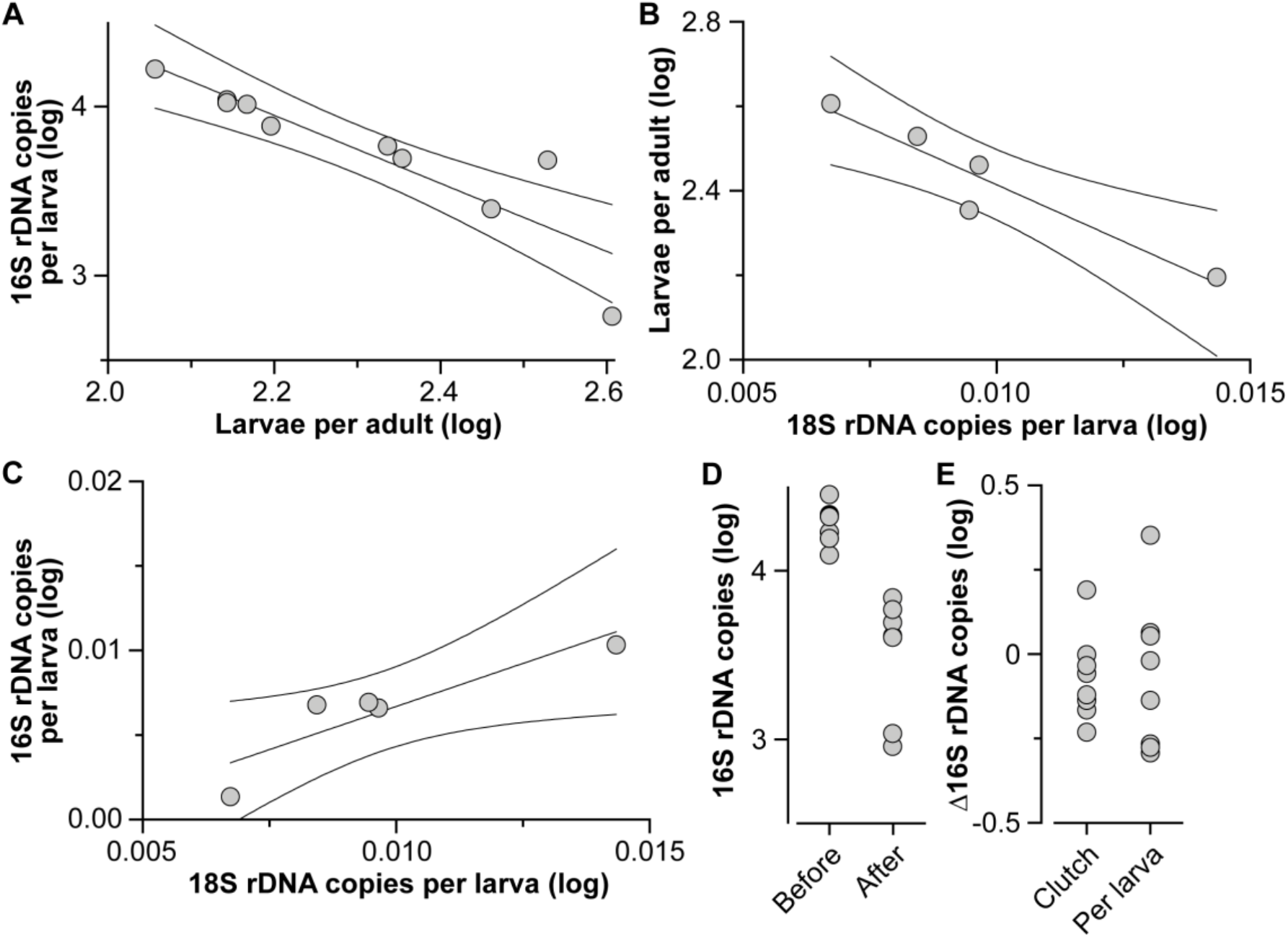
Maternal provisioning of “*Candidatus* Halichondribacter symbioticus.” (A) The reproductive output per adult of *Halichondria panicea* (n = 10) was estimated as the number of larvae per adult and the number of ‘*Ca*. H. symbioticus’ was then quantified from each clutch. The negative correlation indicated a trade-off between fecundity and ‘*Ca*. H. symbioticus’ per larva (F_1,8_ = 32.20, p = 0.0005; R^2^ = 0.801; average ± 95% confidence intervals), such that larvae from more fecund adults had less ‘*Ca*. H. symbioticus’ on average than larvae from less fecund adults. (B) A trade-off was observed between reproductive output and host cells (*i.e*., 18S rDNA copies) per larva (n = 5), such that more fecund adult *H. panicea* tended to produce smaller larvae than less fecund adult larva (F_1,3_ = 27.21, p = 0.014; R^2^ = 0.901; average ± 95% confidence intervals). (C) The smaller larvae from more fecund adult *H. panicea* also had less ‘*Ca*. H. symbioticus’ than the larger larvae from more fecund adults (F_1,3_ = 12.17, p = 0.039; R^2^ = 0.802; average ± 95% confidence intervals). (D) ‘*Ca*. H. symbioticus’ was quantified for adult *H. panicea* (n = 7) before and after spawning and showed a reduction of ‘*Ca*. H. symbioticus’ after reproduction (paired t-test, p < 0.0001). (E) Variation in total ‘*Ca*. H. symbioticus’ between clutches and based on per larva averages. All data types were log transformed for normalization.

A trade-off between reproduction and ‘*Ca*. H. symbioticus’ implies that this obligate symbiont is maternally provisioned [25–27]. Evolutionary theory would then predict that there is a finite maternal symbiont pool that is depleted after spawning and that the abundance this obligate symbiont should not be reduced in the offspring [28–30]. Adults had less ‘*Ca*. H. symbioticus’ after spawning than they did before spawning, and this pattern was also observed for total bacteria (paired t-test, p < 0.0001 for both) (Fig 2D, S2D). Variation in ‘*Ca*. H. symbioticus’ and total bacteria per clutch was, on average, 26.6% and 40.6% of a magnitude, respectively (Fig 2E, S2E). Variation per larva was 57.8% higher for ‘*Ca*. H. symbioticus’ and 17.8% higher for total bacteria than in the adults (Fig 2E, S2E). Furthermore, ‘*Ca*. H. symbioticus’ could not be experimentally reduced in larvae following an antibiotic treatment, while the total bacterial community could be (one-way ANOVA for total bacteria, F = 5.665, p = 0.012; Tukey’s, control vs antibiotics at 48 h: p = 0.01, all other comparisons: p > 0.05; one-way ANOVA for ‘*Ca*. H. symbioticus,’ F = 1.346, p = 0.306) (Fig 3A). A reduction in the total bacteria did not affect the ability of these larvae to settle and undergo metamorphosis (paired t-test, p = 0.137) (Fig 3B).

**Fig 3:**
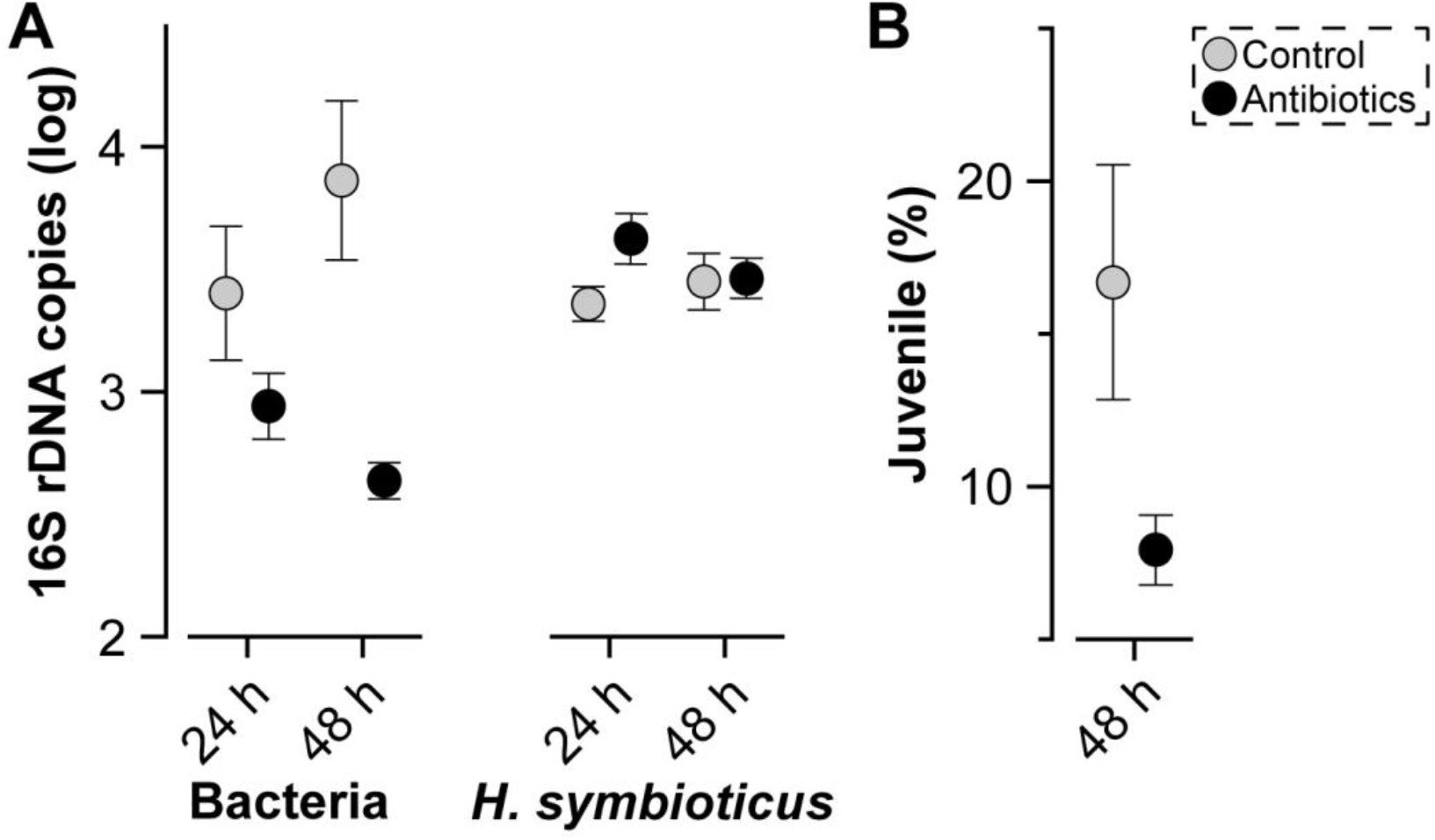
Stability of “*Candidatus* Halichondribacter symbioticus” in larvae. (A) The total bacterial community can be reduced in *Halichondriapanicea* larvae (n = 4, with 50 larvae in each replicate) following 48 h of antibiotics (one-way ANOVA for total bacteria, F = 5.665, p = 0.012; Tukey’s, control vs antibiotics at 48 h: p = 0.01, all other comparisons: p > 0.05), while ‘*Ca*. H. symbioticus’ of could not be reduced (one-way ANOVA, F = 1.346, p = 0.306) (average ± standard error). (B) The reduction in total bacteria did not affect the ability of *H. panicea* larvae to settle and undergo metamorphosis (paired t-test, p = 0.137) (average ± standard error).

If ‘*Ca*. H. symbioticus’ is maternally transmitted and is adaptive for *H. panicea*, then evolutionary theory would predict that there may be multiple strategies of this provisioning [27, 31–33]. We, thus, explored whether the relative abundance of ‘*Ca*. H. symbioticus’ was consistent across individuals. The distribution of the relative abundance for ‘*Ca*. H. symbioticus’ in adults and larvae (on average) each formed two distinct groups: those that have a high or low relative abundance (adults: unpaired t-test, p = 0.0003; larvae: unpaired t-test, p = 0.015) (Fig S3A). These groupings were not always consistent across the life stages for individuals (Fig S3B). Adults with a high relatively abundance of ‘*Ca*. H. symbioticus’ either remained as so (n =3) or switched to a low relative abundance as larvae (n = 4), while adults with a low relative abundance tended to remain that in larvae (n = 3) but rarely switched (n = 1) (Fig S3B, C). The difference in relative abundance between life stages was inconsistent across these ‘provisioning strategies’ (Kruskal-Wallis, H-value = 8.018, p = 0.002), with the only inter-strategy difference being between high-low and low-low (p = 0.015) (Fig S3C).

## DISCUSSION

Vertical transmission can be described as the offspring within a clutch and between clutches being provided a similar number of symbionts irrespective of reproductive output (fecundity) [1–3]. By quantifying the number and size of larvae from *H. panicea* as well as the number of bacteria and ‘*Ca*. H. symbioticus,’ we reject the hypothesis that the offspring of this viviparous sponge are vertically transmitted a similar number of obligate symbionts [5, 8–10]. *H. panicea* appears to take an alternative strategy: a maternal pool of ‘*Ca*. H. symbioticus’ is partitioned proportionally to the reproductive output and allometrically by offspring size. A trade-off between offspring size and number is, by definition, maternal provisioning [28], and parallel differences in ‘*Ca*. H. symbioticus’ abundance to this tradeoff would suggest that this symbiont is also provisioned. Vertically transmitted symbionts can, therefore, be a dynamic process that reflects the reproductive and life-history strategy of the host [1, 2, 34–36].

Mothers are known to influence the development of their offspring through gene transfer and maternal effects [25–27]. These effects are broadly classified and most notably include the resources that are provisioned to an offspring during internal gestation or to the egg of external developers. Typically, these resources include carbohydrates, lipids, mRNAs, and proteins, and the quantity and composition of these resources can have a profound impact on offspring survival, performance, and, thus, fitness [37–40]. Total maternal investment in reproduction, however, is finite and selection balances offspring size and number, such that species tends to either produce many small offspring or few large offspring [28, 41]. As such, energetic content scales with offspring size and this, in turn, increases the likelihood that larger offspring survive [37, 42, 43]. The trade-off between *H. panicea* reproduction and ‘*Ca*. H. symbioticus’ provides the first example of where a vertically transmitted symbiont falls within the framework of maternal provisioning.

The total bacterial community provided to *H. panicea* larvae was inconsistent with this premise. Similar to animal hosts from diverse lineages, the bacterial community associated with sponges is a specific mix of generalists that partially form by neutral processes and a fraction of these microbes are then vertically transmitted [16, 44, 45]. This taxonomic stochasticity has led others to postulate that some maternally transmitted microbes provide little to no fitness to the sponge holobiont, of which could be depleted upon perturbation [11, 16, 46, 47]. An experimental reduction in the total bacterial community that associated with *H. panicea* larvae is consistent with this statement, suggesting that these other bacteria may not have an essential functional role during larval development. Moreover, the inability to perturb ‘*Ca*. H. symbioticus’ in adults [20] as well as larvae implies that this symbiont benefits both maternal and offspring fitness. One potential pattern of this fitness landscape could be that some genotypes have higher or lower relative abundances of ‘*Ca*. H. symbioticus’ and this, in turn, would further influence how mothers influence their offspring. Therefore, the maternal provisioning of ‘*Ca*. H. symbioticus’ is presumed to be adaptive [25–27] and is possibly part of a bet-hedging strategy to maximize maternal fitness in an unpredictable environment, such as that of the Baltic Sea [5, 7, 27, 31, 35, 48–50].

Evolutionary ecology is centered around the examinations of individual life-history characters, of which are increasingly recognized to be dependent on microbial symbionts [51–54]. Provided that holobionts are units of biological organization upon which multilevel selection acts [55, 56], the ways that mothers impact offspring survival, performance, and fitness is anticipated to include the microbes that associate with the offspring [1–3, 5, 35, 48, 57, 58]. Our quantitative estimates of *H. panicea* reproduction and ‘*Ca*. H. symbioticus’ provides the first example of where a vertically transmitted symbiont falls within the framework of maternal provisioning. This raises questions on how widespread this phenomenon is in host-microbe symbioses, what the reproductive and developmental mechanisms are, and how the environment influences the maternal provisioning of obligate symbionts.

## MATERIALS AND METHODS

### Collection of sponge adults and larvae

Adult *H. panicea* were collected at ~2 m depth from the breakwaters at Schilksee Strandbad (Kiel, Germany; 54.424772, 10.175033) during May and June 2020 (n = 30), 2021 (n = 29), and 2022 (n = 14). Sponges were gently removed from boulders using a paint scraper, kept fully submerged in seawater, and transferred to flow-through aquaria at the GEOMAR Helmholtz Centre for Ocean Research (Kiel, Germany) in ~1 hour. Sponges were trimmed to an area of ~45 cm^2^, positioned on terra cotta plates, and mesh traps were placed around each individual [16]. All sponges were provided at least two days to acclimate to the flow-through aquaria.

In 2020, tissues were sampled from each adult before transferring them to aquaria (‘pre-spawn’) as well as after spawning finished (‘post-spawn’). Tissues were preserved in RNAlater at 4°C for 24 hours and stored at −80°C for long-term storage. Sponges were inspected every two days for larvae using a dissecting microscope (Motic, series SMZ168) and everyday thereafter for up to 4 weeks once larvae were observed. Larvae were collected from the bottle (~100 mL) component of each trap as well as by pipetting ~100 mL from the surface off each parental sponge. As has been found previously [59], *H. panicea* larvae were mainly found close to the adult and, thus, only seawater from the surface of each adult was taken in subsequent years. Larvae were preserved in RNAlater at 4°C for 24 h and stored at −80 °C.

In 2021 and 2022, ~100 mL of seawater was pipetted from the surface off each parental sponge on each day for up to 11 days (a total of two weeks in the aquaria when the acclimation period is included). Surface water from each sponge was then transferred to separate jars and the number of larvae were quantified for each adult using a dissecting microscope (Motic, series SMZ168). Two technical replicates of 50 larvae that were pooled from all reproductive adults on that day were transferred to petri dishes and were exposed to either control conditions (0.22 μm filtered sea water) or an antibiotic cocktail (Streptomycin at 25 μg/mL, Penicillin at 10 U/mL, and Rifampicin 30 μg/mL) in early June 2021 (n = 4). One technical replicate was sampled at 24 h while the other was sampled at 48 h. Larvae were collected in sterile 1.5 mL Eppendorf tubes and stored at −80 °C. Juveniles were counted using a dissecting microscope (Motic, series SMZ168).

### DNA extraction and quantitative PCR

Total DNA was extracted from ~90 mg of adult tissue and pooled larvae from each adult according to the manufacture’s protocol for the DNeasy® Blood & Tissue Mini Kit (Qiagen). Total DNA was then quantified using the dsDNA BR Assay Kits for the Qubit Fluorometer (Thermo Fisher Scientific) following the manufacture’s protocol.

Total bacteria was quantified using eubacterial primers for the 16S rRNA gene (F: TGCATGGYTGTCGTCAGCTCG; R: CGTCRTCCCCRCCTTCC; 141 bp) [60], ‘*Ca*. H. symbioticus’ was quantified using specific-specific primers for the 16S rRNA gene (F: CGCGGATGGTAGAGATACCG; R: TGTCCCCAACTGAATGCTGG; 148 bp) [20], and host DNA was quantified using primers for the 18S rRNA gene of Eukaryotes (F: CAGGGTTCGATTCCGTAGAG; R: CCTCCAGTGGATCCTCGTTA; 185 bp) [61]. A standard curve was prepared by running 50 μL PCR reactions as would be prepared for qPCR. Gel electrophoresis was performed on these PCR products, of which were then cut from the gel using a sterile scalpel and cleaned by following the manufacturers protocol in the NucleoSpin Gel and PCR Clean-up kit (Marcherey Nagel). A 10-fold dilution series from 10^−9^ to 10^−3^ ng/μL of DNA was then prepared and the copy number was determined based on the DNA concentration and fragment length.

Each quantitative PCR reaction (totaling 20 μL) was performed in triplicate using 4 μL of template (at 2.5 ng/μL for adult-clutch comparisons and 0.375 ng/μL for all others Fig S4), 10 μL of the Maxima SYBR Green qPCR Master Mix (Thermofisher), 0.1 μL of forward and reverse primers (total of 250 nM), and 5.8 μL of DNAase and RNAase-free water. The following cycle parameters were used on a CFX Connect Real-Time PCR Detection System (Bio-Rad) thermo cycler: 96°C for 2 min, followed by 40 cycles of 94 °C for 30 s, 60 °C for 30 s, and 72 °C for 60 s, and a melting curve analysis was conducted by increasing temperature from 60 °C to 95 °C during 5 s. Absolute copy number was then calculated according to the standard curve.

### Statistics

Quantitative estimates of reproductive output for each adult sponge was normalized by log transformation. Log-transformed data were compared between years using a oneway analysis of variance (ANOVA), a simple linear regression was used for spawning duration and total sponges, and a second order polynomial (quadratic) for day number. A simple linear regression was also used to compare day number and total sponges. Quantitative estimates of bacteria, ‘*Ca*. H. symbioticus’, and host cells (via the 18S rRNA gene) were standardized to 1 ng of DNA and then normalized by log transformation. Log-transformed data were compared between life stages (adults and larvae) using a paired t-test for each data type.

We used a series of simple linear regressions to test whether total bacteria and ‘*Ca*. H. symbioticus’ are maternally provisioning. First, we compared the log-transformed quantifications for total larvae and total bacteria or ‘*Ca*. H. symbioticus’ abundance per larva. Second, we compared the log-transformed quantifications to total larvae as well as the total bacteria or ‘*Ca*. H. symbioticus’ abundance per larva. We then used paired t-tests to determine whether log-transformed quantifications total bacteria and ‘*Ca*. H. symbioticus’ differed for adults before and after spawning. Separate one-way ANOVAs (with Tukey’s pairwise comparisons) were used to test whether total bacteria or ‘*Ca*. H. symbioticus’ were reduced in larvae following an antibiotic treatment, and a paired t-test to determine whether settlement was affected by this experimental exposure. Relative abundance was calculated using the standardized estimates of bacteria and ‘*Ca*. H. symbioticus,’ and the potential transmission strategies were compared using unpaired t-tests. Absolute difference between transmission strategies was compared using a one-way ANOVA; ‘low-high’ was not included here because the replication (n = 1) was insufficient for statistical comparisons.

All analyses were performed in Prism (v. 9.0.0). Graphs were created in Prism and then stylized using Adobe Illustrator.

## Acknowledgements

We thank the staff and Research Unit Marine Symbioses at the GEOMAR Helmholtz Centre for Ocean Research (Kiel, Germany) for logistical assistance with collections and experiments.

## Author Contributions

Conceptualization: Tyler J. Carrier, Lara Schmittmann, Lucía Pita, and Ute Hentschel

Data curation: Tyler J. Carrier

Formal analysis: Tyler J. Carrier

Funding acquisition: Ute Hentschel

Investigation: Tyler J. Carrier, Lara Schmittmann, Sabrina Jung, and Lucía Pita

Methodology: Tyler J. Carrier, Lara Schmittmann, Sabrina Jung, and Lucía Pita

Project administration: Tyler J. Carrier and Ute Hentschel

Resources: Ute Hentschel

Software: Tyler J. Carrier and Ute Hentschel

Supervision: Tyler J. Carrier and Ute Hentschel

Validation: Tyler J. Carrier and Sabrina Jung

Writing – original draft: Tyler J. Carrier

Writing – review & editing: Tyler J. Carrier, Lara Schmittmann, Sabrina Jung, Lucía Pita, and Ute Hentschel

## Supplementary Information

**Fig S1:**
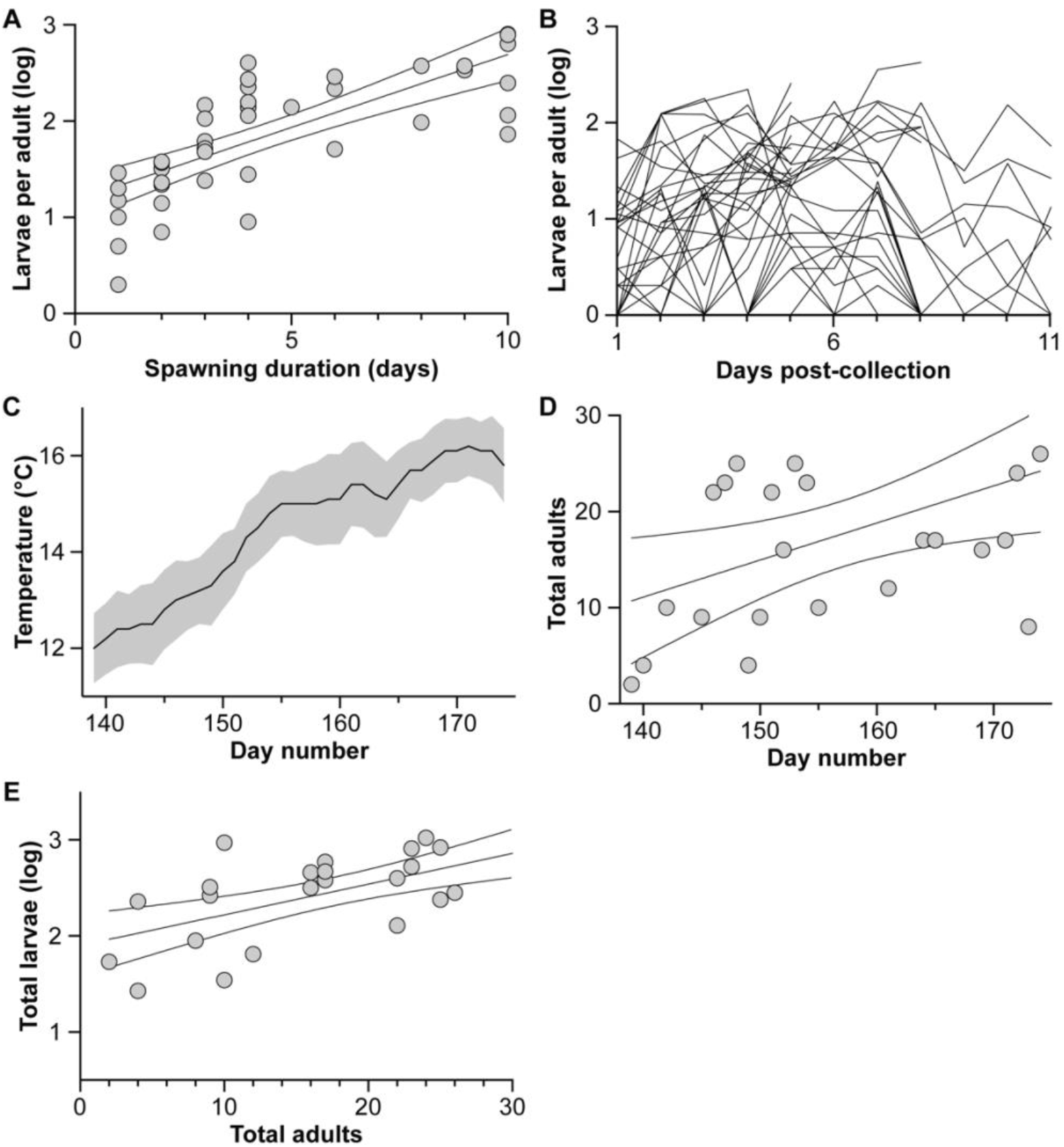
Reproductive output of *Halichondria panicea*. (A) Estimation of the clutch size for individual *H. panicea* and how this output relates to the duration that an individual sponge releases its larvae (F_1,40_ = 50.46, p < 0.0001; R^2^ = 0.558; average ± 95% confidence intervals). (B) The estimated number of released larvae for 54 individuals (*i.e*., all from 2021 and 2022) for up to 11 days after collection. (C) Daily mean temperatures between 19 May and 23 June for the years 1997-2021 (black line: daily mean averages, grey area: 95% confidence intervals) recorded at 1.5 m depth from the Kiel Fjord (54°19’46.0”N, 10°08’58.3”E). These data for 2022 will not be processed and available from the GEOMAR Helmholtz Centre for Ocean Research until mid-2023 and, thus, are not included here. (D) The positive relationship between day-number of the year and total sponges that were being sampled for larvae (F_1,23_ = 6.543, p = 0.018; R^2^ = 0.222; average ± 95% confidence intervals). (E) The positive relationship between the number of sponges that were sampled on a given day and the total larvae that were collected (F_1,23_ = 16.21, p = 0.0005; R^2^ = 0.413; average ± 95% confidence intervals). Most data types were log transformed for normalization.

**Fig S2:**
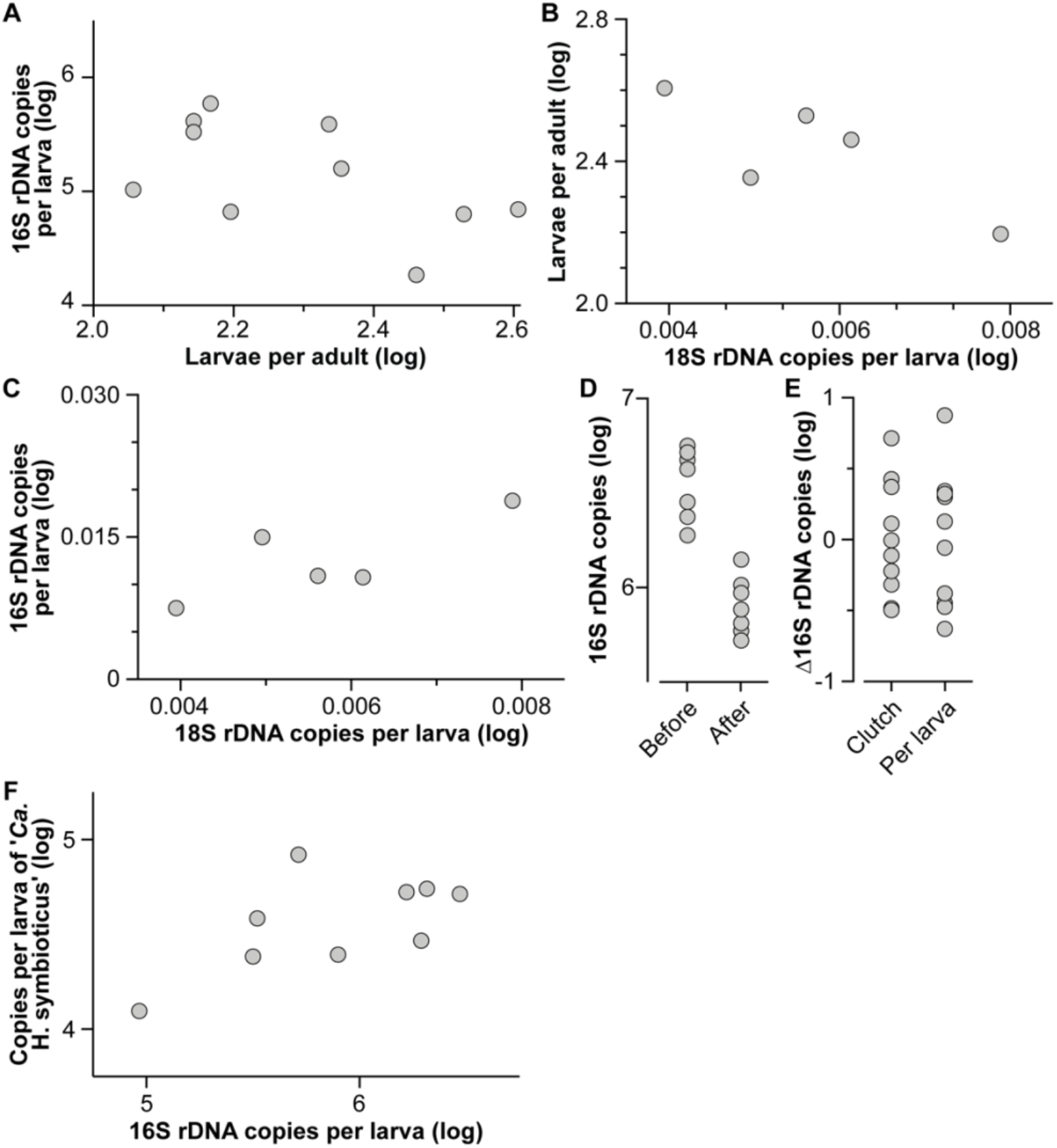
No maternal provisioning of the entire bacterial community. (A) The number of larvae was estimated for adult *Halichondria panicea* (n = 10) and the number of bacteria was then quantified from each clutch. We did not observe a trade-off between fecundity and total bacteria per larva (F_1,8_ = 3.53, p = 0.0971; R^2^ = 0.306). (B) No trade-off was observed between reproductive output and host cells (*i.e*., 18S rDNA copies) per larva (n = 5) (F_1,3_ = 27.21, p = 0.097; R^2^ = 0.656). (C) The smaller larvae from more fecund adult *H. panicea* did not have less bacteria than the larger larvae from more fecund adults (F_1,3_ = 4.74, p = 0.118; R^2^ = 0.612). (D) Total bacteria were quantified for adult *H. panicea* (n = 7) before and after spawning and, consistent with the expectation for maternal provisioning, adults have less bacteria after spawning than they did before spawning (paired t-test, p < 0.0001). (D) Variation in total bacteria between clutches and based on per larva averages. (F) A trend suggestive of a relationship between “*Candidatus* Halichondribacter symbioticus” per larva and total bacteria per larva (F_1,7_ = 4.253, p = 0.078; R^2^ = 0.378). All data types were log transformed for normalization.

**Fig S3:**
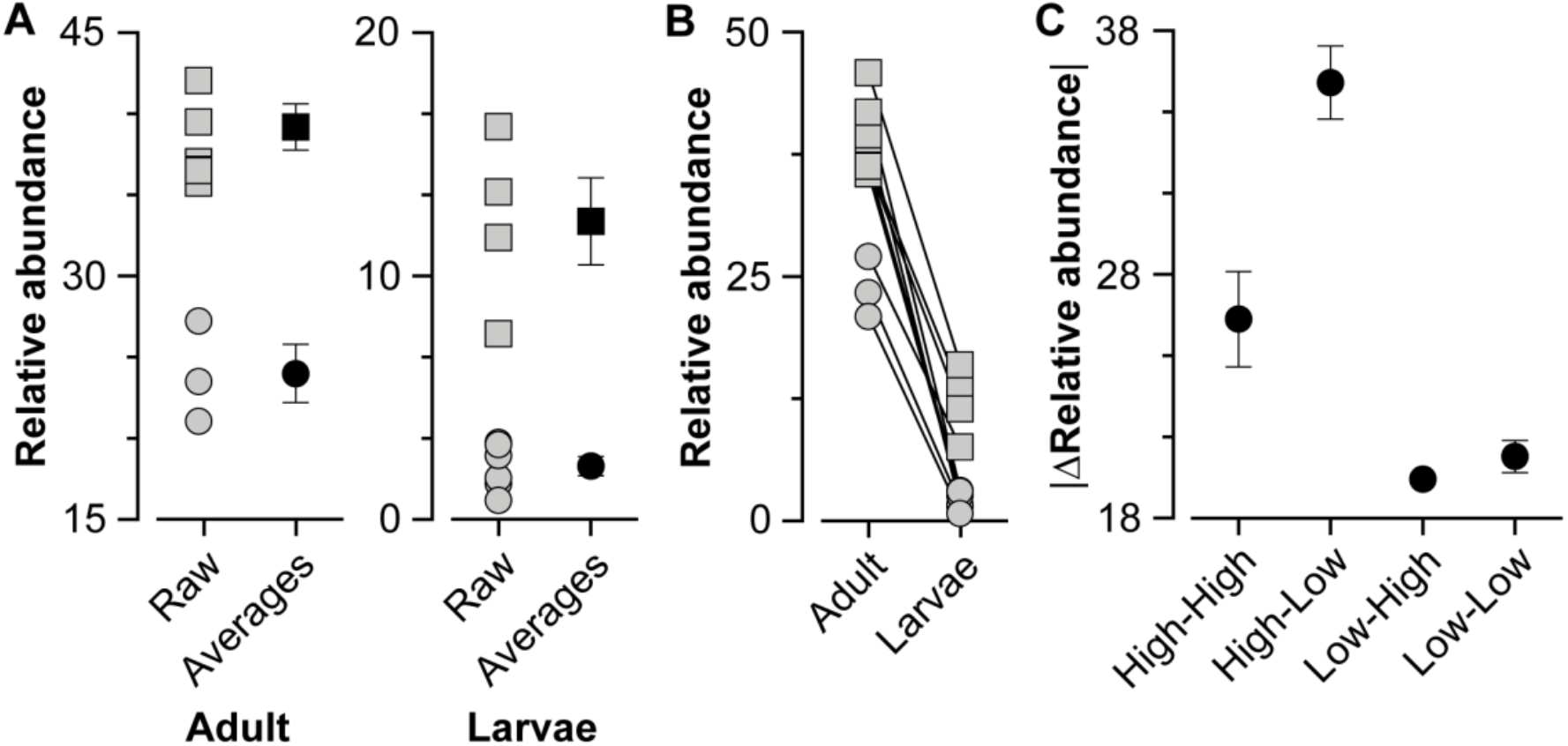
Provisioning strategies of “*Candidates* Halichondribacter symbioticus.” (A) The relative abundance (average ± standard error) of ‘*Ca*. H. symbioticus’ was compared for adults (left) and their larvae (right) (n = 10). Individuals from both life stages formed two distinct group (adults: unpaired t-test, p = 0.0003; larvae: unpaired t-test, p = 0.015), where the abundance of ‘*Ca*. H. symbioticus’ was high (squares) or low (circles) relative to the total bacterial community. (B) Adult *Halichondria panicea* and their larvae that had relatively high and low ‘*Ca*. H. symbioticus’ were paired. (C) Adult-larval pairs that had either a relatively high or low abundance of ‘*Ca*. H. symbioticus’ indicated that individuals may remain relatively high or low for both life stages (n = 3 for each) or be relatively high as adults and relatively low as larvae (n = 4) or vice versa (n = 1). The delta in relative abundance differed between these groups (H-value = 8.018, p = 0.001), with the only inter-strategy difference being between high-low and low-low (p = 0.015).

**Fig S4:**
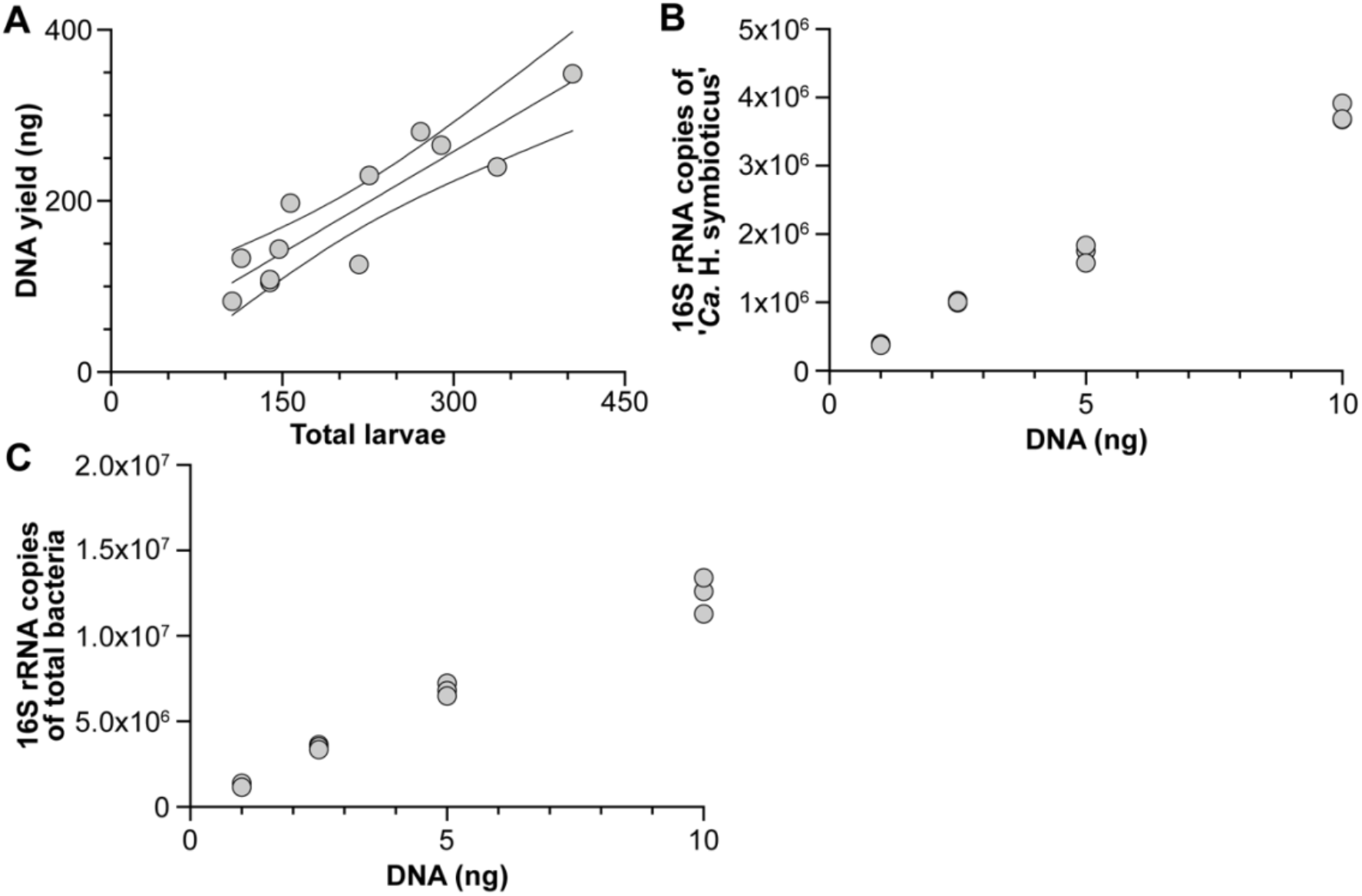
Molecular parameters of *Halichondria panicea* larvae and “*Candidatus* Halichondribacter symbioticus.” (A) The DNA yield increased proportionally with the total number of larvae (F_1,10_ = 41.91, p < 0.0001; R^2^ = 0.807; average ± 95% confidence intervals), such that each larva had on average 0.89 ng of DNA. (B) The detection of bacteria ‘*Ca*. H. symbioticus’ increased proportionally with DNA input and justified using 1.5 ng of total DNA per quantitative PCR reaction were sufficient for reliably detecting and used for all reactions. (C) The detection of total bacteria increased proportionally with DNA input and justified using 1.5 ng of total DNA per quantitative PCR reaction were sufficient for reliably detecting and used for all reactions.

